# Topology-Aware Focal Loss for 3D Image Segmentation

**DOI:** 10.1101/2023.04.21.537860

**Authors:** Andac Demir, Elie Massaad, Bulent Kiziltan

## Abstract

The efficacy of segmentation algorithms is frequently compromised by topological errors like overlapping regions, disrupted connections, and voids. To tackle this problem, we introduce a novel loss function, namely Topology-Aware Focal Loss (TAFL), that incorporates the conventional Focal Loss with a topological constraint term based on the Wasserstein distance between the ground truth and predicted segmentation masks’ persistence diagrams. By enforcing identical topology as the ground truth, the topological constraint can effectively resolve topological errors, while Focal Loss tackles class imbalance. We begin by constructing persistence diagrams from filtered cubical complexes of the ground truth and predicted segmentation masks. We subsequently utilize the Sinkhorn-Knopp algorithm to determine the optimal transport plan between the two persistence diagrams. The resultant transport plan minimizes the cost of transporting mass from one distribution to the other and provides a mapping between the points in the two persistence diagrams. We then compute the Wasserstein distance based on this travel plan to measure the topological dissimilarity between the ground truth and predicted masks. We evaluate our approach by training a 3D U-Net with the MICCAI Brain Tumor Segmentation (BraTS) challenge validation dataset, which requires accurate segmentation of 3D MRI scans that integrate various modalities for the precise identification and tracking of malignant brain tumors. Then, we demonstrate that the quality of segmentation performance is enhanced by regularizing the focal loss through the addition of a topological constraint as a penalty term.

## 1. Introduction

In the domain of image segmentation, several deep learning approaches have been proposed. Notable among them are U-Net [38], Mask R-CNN [19] and SegNet [2]. However, these techniques have limited scalability when applied to large 3D volumes. In order to overcome this limitation, modifications such as multi-scale architectures [45] and attention mechanisms [33] have been suggested. Nevertheless, these approaches still suffer from deficiencies such as limited receptive field, generalization and scalability. Specifically, they can only capture a limited context around the target voxel, which hinders their effectiveness. Although larger kernel sizes or dilated convolutions can address this issue, it comes at the cost of computational complexity. Another challenge is their sensitivity to data variability, which can be addressed through data augmentation and transfer learning. Lastly, the limited scalability of these techniques to larger 3D volumes is attributed to the need for multiple downsampling and upsampling steps that result in information loss and reduced accuracy.

Recent advances in image segmentation using convolutional neural networks (CNNs) have shown remarkable results. However, the performance of these networks can be further enhanced by incorporating topological data analysis (TDA) tools [21,22,32]. TDA tools are useful for analyzing high dimensional data, and can capture important geometric and structural information about the data.

One way to incorporate TDA into CNNs is to add topological layers to the network architecture [28]. These layers can capture the persistence of the homology groups of the input data using a persistence diagram layer. Alternatively, topological features can be computed using TDA methods like persistent homology or mapper, and used as input to the CNN, thereby improving the robustness and accuracy of the network.

In addition, topological loss can be integrated into the training of a neural network to improve its performance for image classification tasks [22, 30]. This loss function measures the topological consistency of the network output with the input data, encouraging the network to learn features that are topologically consistent. This is achieved by modifying the standard cross-entropy loss function to include a term that measures the topological consistency of the network output with the input data using again TDA methods like persistent homology or mapper.

Machine learning methods have advanced to the point where brain tumors can be segmented, and tumors’ genetic and molecular biology can be predicted based on MRI. [39] However, none of the attempted approaches were clinically viable because of limited performance. [6]

Existing methods provide limited capabilities to harness the variability in tumor shape and texture that characterize different types of diffuse gliomas. Hence we propose using topological features computed by persistent homology to identify malignant entities. We hypothesize that topological features contain useful complementary information to image-appearance based features that can improve discriminatory performance of classifiers.

### 1.1. Our contributions

- We introduce a novel loss function, TAFL, that combines the Focal Loss with a topological constraint term based on the Wasserstein distance between the ground truth and predicted segmentation masks’ persistence diagrams.
- We construct persistence diagrams from filtered cubical complexes of the ground truth and predicted segmentation masks and utilize the Sinkhorn-Knopp algorithm to determine the optimal transport plan between the two persistence diagrams.
- We evaluate TAFL by training a 3D U-Net with the MICCAI Brain Tumor Segmentation (BraTS) challenge validation dataset, which requires accurate segmentation of 3D MRI scans for the precise segmentation of malignant brain tumors.
- We demonstrate that the quality of segmentation performance is enhanced by regularizing the focal loss through the addition of a topological constraint as a penalty term. Overall, our approach provides an effective solution to the challenge of topological errors in segmentation algorithms.

## 2. Related Work

### 2.1. Deep Learning in Brain Tumor Detection

Central nervous system (CNS) tumors are among the most fatal cancers in humans of all ages. Gliomas are the most common primary malignancies of the CNS, with varying degrees of aggressiveness, morbidity, and mortality. Currently, most gliomas require tissue sampling for diagnosis, tumor grading, and identification of molecular features to targeted therapies. Accurate segmentation of gliomas is important for surgeons because it can help them better understand the tumor’s location, shape, and size, which can inform treatment planning and improve surgical outcomes.

Upon clinical presentation, patients typically undergo imaging followed by biopsy alone or biopsy with resection for diagnosis and determination of histopathologic classification, tumor grade, and molecular markers. [46] Differen-tiation of low-grade gliomas (LGGs; grade II) from highgrade gliomas (HGGs; grades III, IV) is critical, as the prognosis and thus the therapeutic strategy could differ substantially depending on the grade. [11, 26] Tissue biopsy, while considered the diagnostic gold standard, is invasive and can sometimes be high-risk or inconclusive. The Cancer Genome Atlas (TCGA) report suggests that only 35% of biopsy samples contain sufficient tumor content for appropriate molecular characterization. [43] Hence, robust noninvasive approaches are desired for accurate diagnosis and monitoring of tumor evolution.

The state-of-the-art models in brain tumor segmentation are based on the encoder-decoder architectures, with U-Netlike architectures [40] (basic U-Net [14, 38], UNETR [18], Residual U-Net [25], and Attention U-Net [33]), were among top submissions to the BraTS challenge. Among factors influencing models performance like architecture modifications, training schedule, and integration of multiple MRI sequences, the size of brain tumor datasets remains a major limiting factor. Additionally, based on earlier successes for classification of tumor grades (high grade vs. low grade), a few studies explored the determination of glioma mutation status (O6-Methylguanine-DNA methyltransferase (MGMT) promoter methylation status) using radio-genomics features. Of importance, analysis of MGMT promoter methylation is essential to predict prognosis and treatment response. As a result, TDA has the potential to support future research in MRI-based deep learning to reveal clinically significant tumor features and aid clinical decision making.

According to several ablation studies run by [14] in order to select the most optimal CNN architecture for 3D medical image segmentation, UNet [38] attains better tumor segmentation performance as measured by Dice score compared to Attention UNet [33], Residual UNet [20], SegResNetVAE [31] and UNetR [18]. Hence, we adopt the U-Net architecture to encode each MRI scan image into a latent-space representation.

### 2.2. TDA in Computer Vision

Persistent homology, which is the primary tool in topological data analysis (TDA), has demonstrated remarkable efficacy in pattern recognition for image and shape analysis. Over the past two decades, numerous works in a variety of fields have utilized persistent homology for a range of applications, including the analysis of hepatic lesions in images [1], human and monkey fibrin images [7], tumor classification [12,35,36], fingerprint classification [16], diabetic retinopathy analysis [13,15], the analysis of 3D shapes [42], neuronal morphology [24], brain artery trees [5], fMRI data [37, 44], and genomic data [9]. Of note, the TDA Applications Library [17] catalogs hundreds of compelling applications of TDA in various fields. For a comprehensive survey of TDA methods in biomedicine, see the survey [41].

On the other hand, TDA has also been successfully applied to the problem of tumor classification in several works, such as [12, 35, 36]. In [36], the authors employed TDA techniques to study histopathological images of colon cancer and obtained impressive results. In [35], the authors studied hepatic tumors using MRI images of the liver, using TDA as a feature extraction method from 2D MRI slices. In [12], the authors examined brain cancer by utilizing a sequence of binary images obtained via tumor segmentation, and capturing topological patterns of brain tumors through the Euler Characteristics Curve in different radial directions.

Although these methods use different variations of sublevel filtrations, they all utilize 2D images and thus differ substantially from our approach. 2D images are limited to 0 and 1-dimensional topological features (components and loops), relying solely on these lower-dimensional features for their study. In contrast, our approach employs 3D MRI images (volumes), with key features derived from 2-dimensional topological features that represent cavities in 3*D* cubical complexes.

#### 3. Background

### 3.1. Persistent Homology

Persistent homology (PH) studies how the topology of a space changes as we vary a parameter that controls the level of detail or resolution of the space. It is particularly useful for studying data sets that have a natural ordering or hierarchy. The main difference between homology and PH is that homology provides a static snapshot of the topology of a space, while PH captures the evolution of the topology over different scales or levels of detail. In other words, homology is useful for understanding the global structure of a space, while PH is better suited for detecting and characterizing the local features of a space.

The core concept behind PH is to describe the changes in homology that occur as an object evolves with respect to a parameter. Given a simplicial complex, Σ, we produce a finite sequence of subcomplexes: Σ_*f*_ = {Σ_*p*_|0 ≤ *p* ≤*m*} such that ∅= Σ_0_ ⊆Σ_1_ ⋯ ⊆Σ_*m*_ = Σ using a filtration technique. Figure 1 illustrates the evolution of the homology groups over the entire sequence of simplicial complexes.

**Figure 1.**
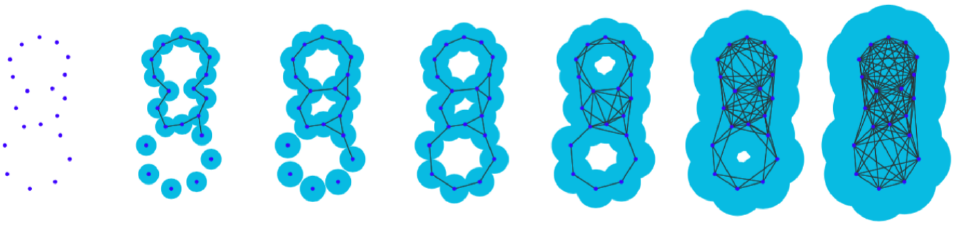
The filtration procedure begins with a set of points and proceeds by incrementally increasing the radius of the balls centered on each point. This procedure constructs a sequence of simplicial complexes that capture the topological features of the space in increasing levels of detail or resolution. Image is from [8].

### 3.2. Cubical Complex Filtration

In this paper, we present a method for applying PH to 3D MRI images, which involves inducing a nested sequence of 3D cubical complexes and tracking their topological features. We represent a 3D MRI image 𝒳 as a 3D cubical complex with resolution *r* × *s* × *t*. To perform PH, we first create a nested sequence of 3D binary images by considering grayscale values *γ*_*ijk*_ ∈ [0, 255] of each voxel Δ_*ijk*_ ⊂𝒳. For a sequence of grayscale values (0 ≤*t*_1_ *< t*_2_ *< ⋯ < t*_*N*_ ≤255), we obtain 3D binary images 𝒳_1_ ⊂𝒳_2_ ⊂ ⋯ ⊂𝒳_*N*_, where 𝒳_*n*_ is the set of voxels Δ_*ijk*_ ⊂𝒳 with *γ*_*ijk*_ ≤*t*_*n*_. In other words, we start with an empty 3D image of size *r* × *s* × *t*, and activate voxels by coloring them black as their grayscale values exceed the threshold, *t*_*n*_, at each step. This process is illustrated in Figure 2. The PH algorithm then tracks the topological features, such as connected components, holes/loops, and cavities, across this sequence of binary images [10, 23]. Overall, this method allows us to gain insights into the complex topological structures of 3D MRI images and can be used for various applications, such as medical image classification and segmentation.

**Figure 2.**
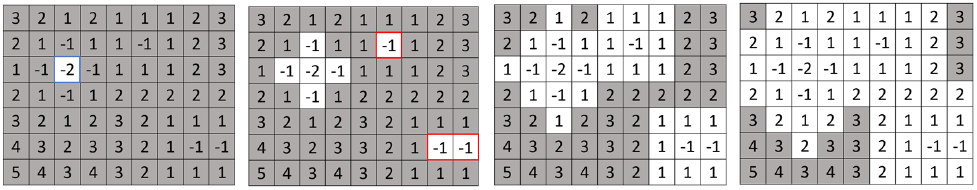
At *t*_*n*_ ≤ −2, a connected component is born, and at *t*_*n*_ ≤ −1, two other connected components are born. When *t*_*n*_ ≤1, the upper component with birth-time −1 is killed and merged into the one with birth-time − 2. Finally, when *t*_*n*_ ≤2, the right component with birth-time −1 is killed. As a result, the 0-th barcodes of the dataset can be computed and represented as [− 2, inf), [− 1, 1), and [− 1, 2). This barcode representation provides a concise and informative summary of the topological structure of the dataset. Image is from [23].

PH tracks the evolution of topological features across a sequence of cubical complexes 𝒳_*n*_ and records it as a persistence diagram (PD). The PD provides information on the birth and death times of topological features, such as connected components, loops, and voids, which appear and disappear over the sequence. Specifically, if a topological feature *σ* first appears in 𝒳_*m*_ and disappears in 𝒳_*n*_, we refer to *b*_*σ*_ = *t*_*m*_ as the birth time and *d*_*σ*_ = *t*_*n*_ as the death time of the feature *σ*. The PD for dimension *k* is then the collection of all such 2-tuples *PD*_*k*_(𝒳) = (*b*_*σ*_, *d*_*σ*_), which represents the persistence of the *k*-dimensional topological features across the sequence. This approach enables us to identify the significant topological features in the data and analyze their persistence over different scales or levels of resolution.

## 4. Measuring Pairwise Distances Between Persistence Diagrams

### 4.1. Sinkhorn-Knopp Algorithm

The optimal transport problem seeks to find the most efficient way to transport a set of masses from one distribution to another, where the cost of transporting a mass depends on the distance between its source and destination points. The Wasserstein distance is not explicitly used in the equations of Sinkhorn’s algorithm, but it arises naturally from the optimization problem being solved.

In the context of persistence diagrams, the masses correspond to the persistence pairs, and the cost of transporting a pair depends on the distance between its birth and death values. The cost matrix *C* used in Sinkhorn’s algorithm encodes these distances, and the optimal transport plan *P* determines how to match the pairs in the two diagrams in order to minimize the total cost. Thus, the Wasserstein distance arises implicitly from the optimal transport problem being solved by Sinkhorn-Knopp algorithm. The final output of the algorithm (the optimal transport plan *P*) can be used to compute the Wasserstein distance between the persistence diagrams.

#### Definition 4.1.

Doubly stochastic matrix, denoted by *D* ∈ ℛ^*N ×N*^, is a square matrix with non-negative entries that is both row-normalized and column-normalized:

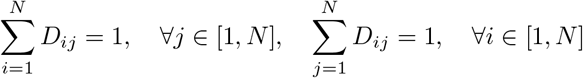

#### Definition 4.2.

The Sinkhorn-Knopp Algorithm states that any square matrix comprising strictly positive elements can be transformed into ∧_1_*D* ∧_2_, where ∧_1_ and ∧_2_ are diagonal matrices with strictly positive diagonal entries.

Sinkhorn-Knopp Algorithm approximates the doubly stochastic matrix with linear convergence. The algorithm operates by iteratively normalizing the matrix along its rows and columns:

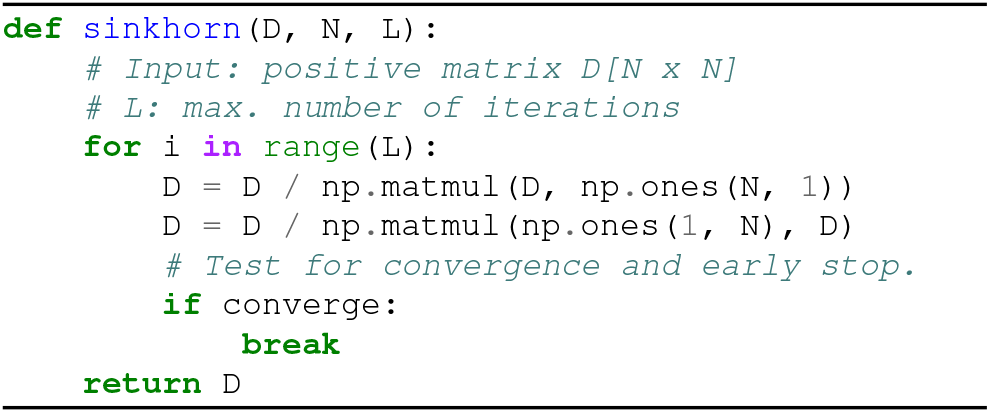

### 4.2. Optimal Transport Plan

#### Algorithm 1

Sinkhorn’s algorithm for optimal transport

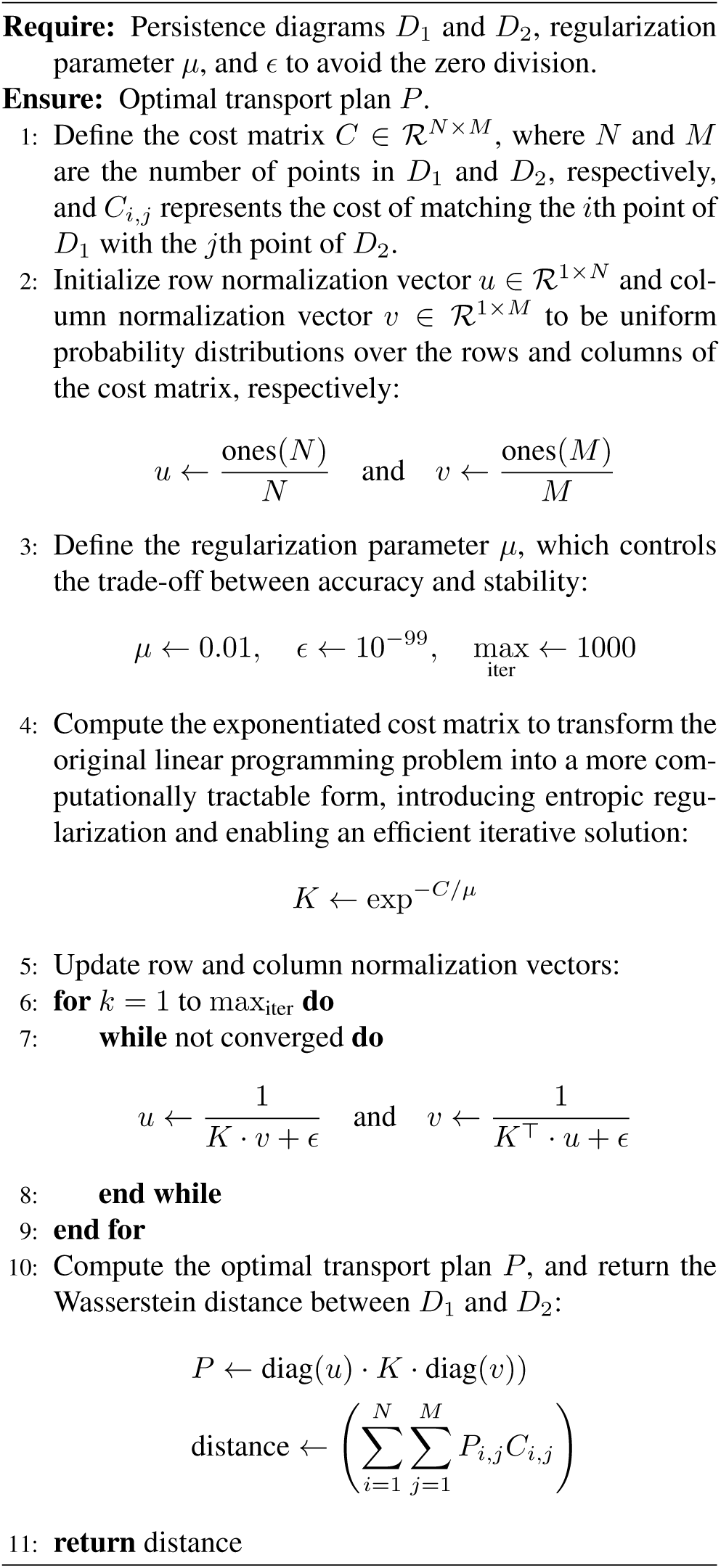

The resulting optimal transport plan *P* represents the most efficient way to match the points of *D*_1_ to the points of *D*_2_. This algorithm is used in a variety of applications, including shape matching and image registration. Next, we present a Python code implementation for computing the optimal transport plan between two persistence diagrams utilizing the Sinkhorn-Knopp algorithm and produce the Wasserstein distance with an example.

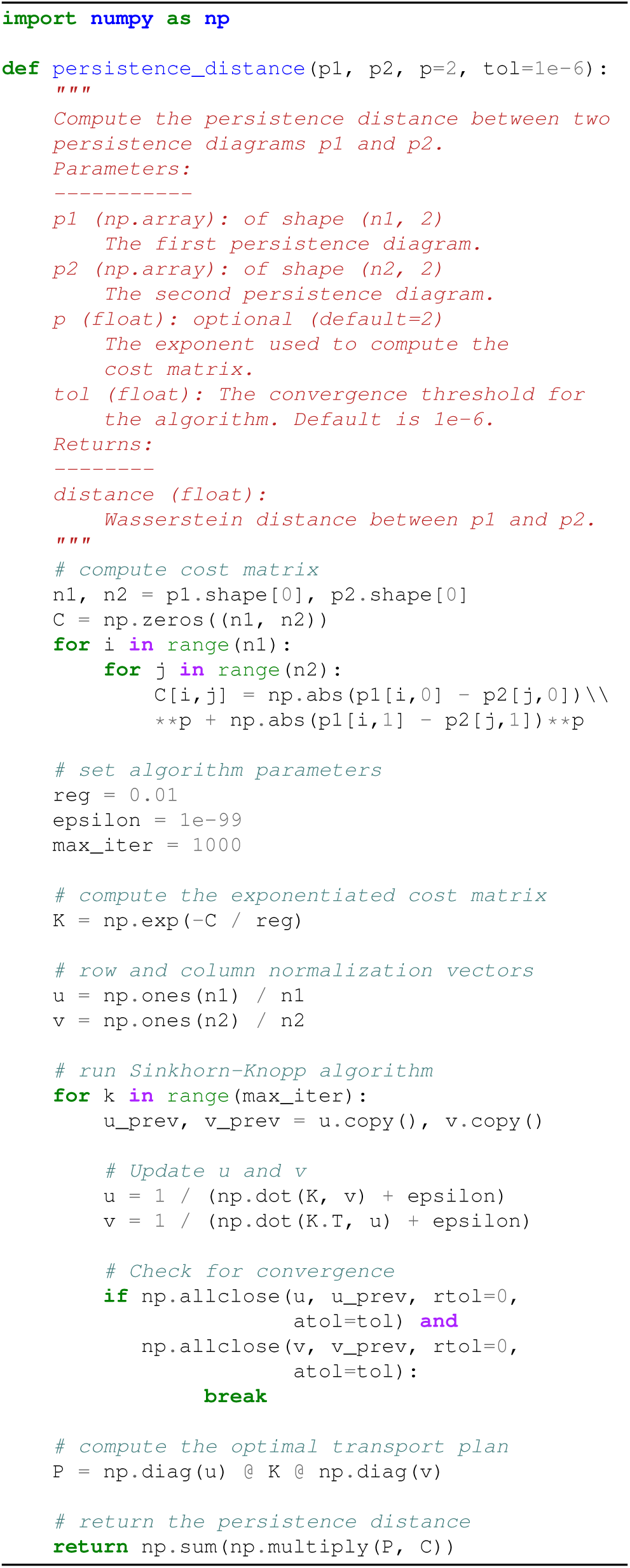

We can provide an illustrative example to demonstrate the efficacy of our approach. Let’s define 2 persistence diagrams as a NumPy array, where the first column corresponds to the birth time and the second column denotes the death time of a topological feature,

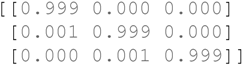

We obtain the resultant cost matrix, *C* as follows,

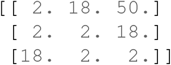

The optimal transport plan matrix, *P* ∈ ℛ^*N×M*^ represents the optimal way to match the points in the two input persistence diagrams, *p*_1_ and *p*_2_, in order to minimize the cost of the matching. Each element *P*_*i,j*_ of the matrix represents the amount of mass that should be transported from point *i* in *p*_1_ to point *j* in *p*_2_, in order to minimize the overall cost of the matching. *P* is subject to row-wise and column-wise normalization. These constraints maintain the conservation of mass throughout the matching process, as they ensure that the total mass remains constant in each row and column.

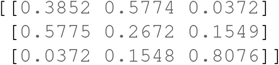

Ultimately, the computed Wasserstein distance yields a value of 6.0.

### 4.3. Topology-Aware Focal Loss

Our approach attains per-pixel precision as well as topological correctness through the utilization of topology-aware focal loss (TAFL), *L*_*tafl*_(*f, g*) by training a 3D U-Net for segmentation. *f* denotes the predicted segmentation mask, while *g* represents the ground truth. The training loss consists of a weighted combination of per-pixel focal loss, *L*_*focal*_ and the aforementioned topological loss, *L*_*topo*_ that is the elementwise sum over the Hadamard product of optimal transport plan matrix, *P*, and cost matrix, *C*.

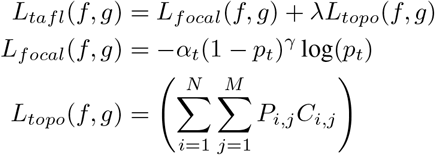

where *λ* is the regularization parameter that modulates the weight of topological loss. In our experiments, we have found that setting *λ* equal to 0.001 yield optimal stability in loss convergence. Additionally, *α* is the weighting factor for focal loss and set to 1 and *γ* is the scaling factor that addresses the issue of class imbalance in image segmentation tasks, whereby well-classified samples, *p*_*t*_ *>* 0.5, are given a lower loss while misclassified samples are emphasized more. We have set the value of *γ* to 2 in our experiments. The probability estimate for each class is denoted by *p*_*t*_.

## 5. Experiments

### 5.1. Datasets

**BraTS 2019** dataset^1^ consists of 335 3D brain MRI scans, segmented manually by one to four experienced neuroradiologists ensuring consistency in the tumor annotation protocol [3, 4, 27, 29]. 259 of them belong to patients diagnosed with HGG and 76 of them to those diagnosed with LGG. Segmentation annotations comprise four classes: the necrotic and non-enhancing tumor core (NCR/NET–label 1), the peritumoral edema (ED–label 2), background (voxels that are not part of the tumor–label 3) and GD-enhancing tumor (ET–label 4). Each MRI scan image has 4 modalities: native (T1) and post-contrast T1-weighted (T1Gd), T2-weighted (T2), and d) T2 Fluid Attenuated Inversion Recovery (T2-FLAIR) and each modality has dimensions of (240, 240, 155) voxels. The provided data is skull-stripped and interpolated to the same resolution of 1 mm^3^.

### 5.2. Preprocessing

Data augmentation transforms were applied to the batches of data sampled from training set to increase the robustness of the U-Net segmentation model and learning a higher quality latent-space representation. In order to mitigate the overfitting problem while training U-Net for tumor segmentation, we synthetically increase the size of the training set by a composition of transforms, which includes: random cropping, rotating, injecting Gaussian noise, adding Gaussian blur, changing brightness and contrast. Specifi-cally, we randomly crop patches of size (5, 128, 128, 128) per MRI scan id, where 5 denotes 4 MRI modalities and their corresponding segmentation mask. Then we apply random flipping along the x, y and z axes with a probability of 0.5. This is followed by injecting Gaussian noise (with mean, *μ* = 0 and standard deviation, *σ* ∼ 𝒰 [0, 0.33]) with a probability of 0.2 and adding Gaussian blur (where *σ* of the Gaussian kernel is sampled from a uniform distribution: *σ* ∼ 𝒰 [0.5, 1.5] with a probability of 0.2. We jitter brightness with a brightness factor uniformly sampled from 𝒰[0.7, 1.4] and contrast with a contrast factor uniformly sampled from 𝒰[0.7, 1.4] at probability of 0.2.

Let *f* (*x*) denote a function approximator, which is a CNN, and *g*(*x*) an image transform, specifically translation that shifts a voxel in the 3D volume such that *I*(*x, y, z*) = *I*(*x* − a, y − b, z − c), where *I*(*x, y, z*) denotes the voxels in the 3D image and *a, b, c* are scalars. *f* (*x*) is said to be equivariant with respect to *g*(*x*), if *f* (*g*(*x*)) = *g*(*f* (*x*)). CNNs are inherently equivariant with respect to translation thanks to the concept of weight sharing, however they are not naturally equivariant to other transformations such as changes in scale, rotation or perturbation with noise. Utilizing translational equivariance by applying transformations such as flipping, zooming or translating in conjunction with noise perturbation and color jittering on each batch of data during training phase comes handy to increase the performance of CNNs and make them more robust against adversaries. However, data augmentation transforms are not expected to achieve the desired outcome while producing persistent homology features, since persistent homology already generates topological features that persist across several scales. An important property of topological features is that they are homotopy invariant as they do not change, when the underlying space is transformed by stretching, bending or other deformations [34]. Hence, we exempt the training data from augmentation transforms while operating cubical persistent homology to ease the computational complexity.

### 5.3. Macro Design

Figure 3 shows that the U-Net architecture is composed of 2 parts, i.e., encoder and decoder. Encoder has 7 convolutional blocks, where each block consists of 2 repeating convolutional layers. The first layer applies Conv3d kernels with size (3 *×* 3 *×* 3) and stride (2, 2, 2), followed by InstanceNorm3d and GeLU activation, whereas the second layer applies kernels of size (3 *×* 3 *×* 3) and stride (1, 1, 1), also followed by InstanceNorm3d and GeLU. We adopt a “patchify” strategy, and eliminate the pooling layer by applying convolutional kernels with stride (2, 2, 2), which effectively yields a (2, 2, 2) downsampling while preserving shift-equivariance. Furthermore, we replace convex and monotonic ReLU activation with a non-convex and non-monotonic activation function GeLU, which is differentiable in its entire domain and allows non-zero gradients for negative inputs unlike ReLU.

**Figure 3.**
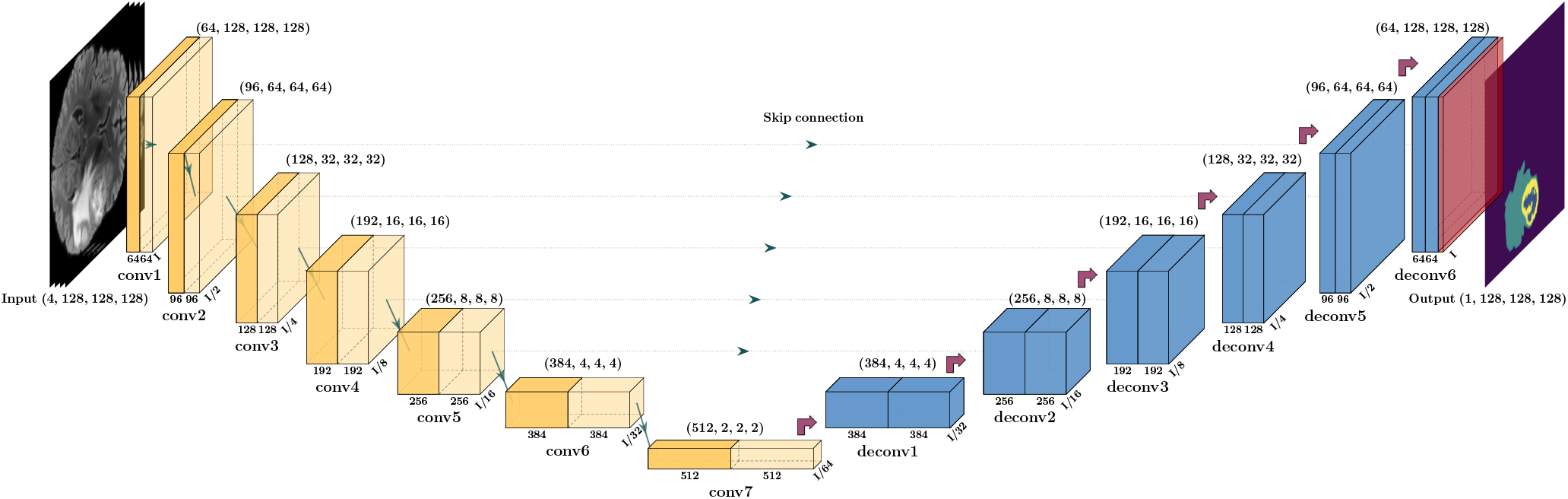
3D U-Net Architecture. The U-Net design employs an encoder module that applies spatial dimension reduction to the input, which is subsequently upsampled by the decoder module to restore the input to its original shape.

After generating a latent vector, ConvTranspose3d operators of kernel size (2 *×* 2 *×* 2) and stride (2, 2, 2), represented with purple arrows in Figure 3, increases the spatial dimensions of the output from previous layer by a factor of 2 until the spatial dimensions of the decoder’s output is equivalent to the original input image. An upsampled feature map is concatenated with the encoder feature map of the equivalent spatial level (except of those between conv6 and deconv1 blocks). Symmetrical long skip connections between encoder and decoder blocks, represented with green, dotted lines, enhances the reconstruction of finegrained details. As far as we know there is no previous research showing the theoretical justification of why long skip connections improve the reconstruction performance, but we conjecture that these lengthy skip connections utilized to transmit the feature maps from early encoder layers to late decoder layers allow to recover the spatial information that is lost during downsampling. The output feature map is passed to a decoder block which consists of 2 repeating, identical, Conv3d layers with kernel size (3 *×* 3 *×* 3) and stride (1, 1, 1). Similary, each Conv3d layer is followed by InstanceNorm3d and GeLU.

To enhance the robustness of our predictions, a well-established approach involves the implementation of test time augmentations. Specifically, during the inference process, we generate multiple variants of the input volume by applying one of eight potential flips along the x, y, and z axes. For each of these versions, we perform inference and subsequently transform the resulting predictions back to the original orientation of the input volume by utilizing the same flips as those employed during input volume processing. Ultimately, we aggregate the probabilities derived from all predictions by computing their average.

### 5.4. Training Schedule

Our approach involves training models from scratch, with no use of fine-tuning. We do not utilize any preexisting models as a starting point and initialize the weights of our model randomly. This allows us to fully explore the model’s parameter space, without the bias or constraints imposed by a pre-existing model. We utilized the Adam optimizer with a learning rate of 0.0005 and a weight decay set to 0.0001 for 200 epochs. To ensure effective training, we implemented a cosine annealing scheduler to dynamically adjust the learning rate throughout the training process. We saved the checkpoint with the highest mean Dice score achieved on the validation set during training.

### 5.5. Empirical Evaluations

ET Dice score quantifies the extent of intersection between the ground truth and the segmented enhancing tumor region. WT amalgamates the edema, enhancing tumor and necrotic tumor core classes to evaluate the degree of overlap. Finally, TC employs the segmented necrotic tumor to evaluate the degree of similarity between the ground truth and the predicted segmentations. The experimental results presented in Table 1 show that training with topology-aware focal loss outperforms training with cross-entropy and focal loss. To ensure fair comparison, we follow a standardized evalution process by using the same training and validation dataset, same U-Net architecture and epoch size. Please note that in the context of brain tumor segmentation, a good Dice score would depend on various factors such as the complexity of the tumor structure, the size and shape of the tumors, and the quality of the input data. Generally, a Dice score of 0.8 or higher is considered to be a good performance for brain tumor segmentation. See Figure 4 for the segmentation results recorded on the validation set.

**Table 1.** The average Dice scores for the ET, TC and WT classes, as well as for all classes combined.

| Loss Objective | ET Dice | TC Dice | WT Dice | Dice |
| --- | --- | --- | --- | --- |
| Cross-Entropy | 0.7255 | 0.7240 | 0.8479 | 0.9570 |
| Focal | 0.7244 | 0.7405 | 0.8601 | 0.9582 |
| **T AFL** | **0.7383** | **0.8005** | **0.8793** | **0.9602** |

**Figure 4.**
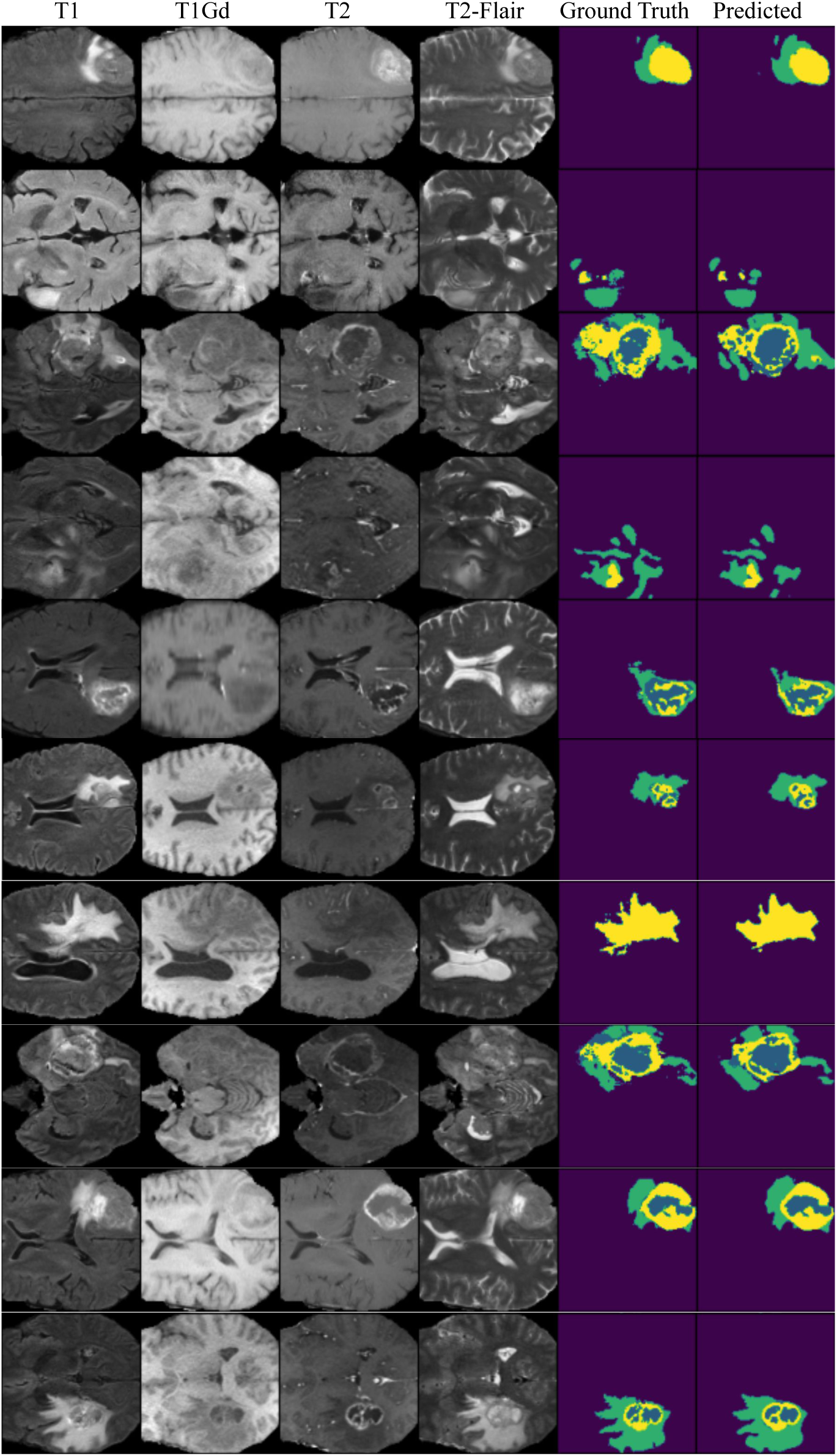
The visualization of segmentation outcomes involves the display of a 3D brain MRI scan image slice featuring modalities T1, T1Gd, T2, and T2-Flair, as well as both ground truth and predicted segmentation masks. To aid in interpretation, color coding is utilized, with purple representing the background or healthy cells, green representing peritumoral edema, yellow indicating an enhancing tumor, and blue representing the necrotic and non-enhancing tumor core.

## 6. Conclusion

We have introduced a novel loss function to address topological errors that frequently arise in segmentation algorithms. We evaluated TAFL on the BraTS challenge validation dataset, and the results showed that it enhances the quality of segmentation performance. Our future work involves investigating the generalization of TAFL to other segmentation tasks and extending its applicability to a wider range of medical imaging modalities.

## Footnotes

1 https://www.med.upenn.edu/cbica/brats2019/data.html

